# Co-culture of type I and type II pneumocytes as a model of alveolar epithelium

**DOI:** 10.1101/2021.03.08.434368

**Authors:** Oliver Brookes, Sonja Boland, René Lai Kuen, Armelle Baeza-Squiban

## Abstract

The epithelial tissues of the distal lung are continuously exposed to inhaled air, and are of research interest in studying respiratory exposure to both hazardous and therapeutic materials. Pharmaco-toxicological research depends on the development of sophisticated models of the alveolar epithelium, which better represent the different cell types present in the native lung and interactions between them.

We developed an air-liquid interface (ALI) model of the alveolar epithelium which incorporates cell lines representative of both type I (NCI-H441) and type II (hAELVi) epithelial cells. We compared morphology of single cells and the structure of cell layers of the two lines using light and electron microscopy. Working both in monotypic cultures and cocultures, we measured barrier function by trans-epithelial electrical resistance (TEER), and demonstrated that barrier properties can be maintained for 30 days. We created a mathematical model of TEER development over time based on these data in order to make inferences about the interactions occurring in these culture systems. We assessed expression of a panel of relevant genes that play important roles in barrier function and differentiation.

The coculture model was observed to form a stable barrier akin to that seen in hAELVi, while expressing surfactant protein C, and having a profile of expression of claudins and aquaporins appropriate for the distal lung. We described cavities which arise within stratified cell layers in NCI-H441 and cocultured cells, and present evidence that these cavities represent an aberrant apical surface. In summary, our results support the coculture of these two cell lines to produce a model which better represents the breadth of functions seen in native alveolar epithelium.

## Introduction

The alveoli of the human lung are the site of all gas exchange required to support survival of the individual. The interface between the blood and breathing gases consists of a composite epithelium comprised of multiple cell types. The epithelium itself features a heterogeneous population of flattened (Type I) and columnar secretory (Type II) cells. The epithelial layer grows in direct contact with the endothelial cells of the underlying capillary circulation; and the compartment features stromal fibroblasts within the alveolar septa and macrophages which enter the tissue from the bloodstream. This complex milieu plays many important roles in supporting breathing, gas exchange and related faculties within the lung. Numerous conditions exist that impair lung function (fibrosis, lung cancer, asthma) which continue to be highly important topics for medical research and create demand for the development and testing of new therapeutic substances, especially using nanomaterials. Moreover, inhalation represents a major route of exposure to airborne toxins, particulates and pathogens, the tissues lining the respiratory tract being at once the principal site of exposure and a major route via which systemic exposure is likely to occur. As a result, in vitro models which accurately and reproducibly represent human lung tissues are highly desirable (1). These models would ideally reflect the cellular composition, structure and signalling interactions seen in the native tissue, as well as recreating functions such as paracellular barrier formation, surfactant production, water homeostasis and protein clearance, which are required to fulfil the role of the tissue in vivo even though they do not directly mediate gas exchange.

The interfacial location of the alveolar epithelium also presents distinct environmental challenges, with multiple distinct adaptations having evolved to support survival and function of the tissue. Therefore, cellular models need to reflect not only the impact of toxins, treatments and nanomaterials on the health of individual cell populations, but also the effect on cellular functions and interactions which permit function on level of the tissue or whole organ. Animal models are widely used but have a high cost in terms of time, expense, and animal welfare. Furthermore, exhaustive hazard assessment with animal models is often impractical and may misrepresent hazards specific to human physiology. There is therefore a drive to create more complex *in vitro* models of tissues which recapitulate elements of the structure and function of the tissue to predict the impact of specific treatments more comprehensively at the tissue level. Numerous existing *in vitro* models use primary cells or immortalised cell lines growing on a porous plastic membrane at the air-liquid interface (2). The plastic membrane system is robust, versatile and has been extensively commercialised in the form of inserts that fit into standard multi-well plates. This paradigm is therefore particularly well suited to standardised tests for assessing toxicological hazards or screening panels of prospective substances of therapeutic interest. While primary cells are more closely representative of the full in-vivo phenotype, they exhibit a high degree of variability between donors, are subject to phenotypic change over time in culture and are not readily available to all researchers. Immortalised cell lines, by contrast, are widely commercially available and produce results which are generally consistent between experimental iterations. Nonetheless, these practical and cost-effective models have their limitations and in order to be able to use this design paradigm, it is necessary to characterise the system with respect to the parameters of the investigation. In the case of distal lung models, this translates to determining whether an adequate paracellular barrier is formed, whether fluid clearance occurs and whether surfactant is produced.

Recent advances in this type of model have demonstrated interactions between cell lines representative of epithelial cells and those representative of the endothelial cells of the pulmonary microvasculature (3, 4) as well as macrophages which typically inhabit the alveolar lumen (5, 6). As the alveolar epithelium consists of two distinct types of epithelial cell: type I cells which line most of the inner surface of the alveoli in close contact with underlying capillary endothelial cells and are flattened to facilitate gas exchange, and type II cells which produce surfactant and act as a progenitor cell population for type I cells. As these cell types are known to interact to inform cell behaviours central to the function of the native tissue, we sought to create a coculture model using cells which are representative of Type I and Type II epithelial cells. NCI-H441 is an adenocarcinoma cell line that is thought to be derived from type 2 epithelial cells, on the strength of its ability to form tight junctions and expression of type 2 markers such as surfactant proteins. These cells have been used to represent the type 2 epithelial cells since their description in the 1980s (7). By contrast, hAELVi is a recently created, line derived from primary type 1 alveolar epithelial cells immortalised using a novel lentiviral transfection method (5, 8). Monolayers of hAELVi cells can form tight-junctional epithelia which can be maintained for long periods, potentially allowing chronic testing of toxicological or therapeutic samples.

Here we describe and compare several structural and functional characteristics of simple models comprising hAELVi or NCI-H441 maintained in culture at the air-liquid interface. We go on to describe a co-culture model consisting of an equal mixture of the two cell lines, present quantitative measurements of the morphology of the epithelium, and describe an unusual artefact of this culture regime.

## Methods and Materials

### Cell culture

NCI-H441 cells (7) were purchased from ATCC (HTB-174). hAELVi cells (8) were purchased from Inscreenex (Brunswick, Germany).

Advanced DMEM (Gibco, France, 12491-015, aDMEM hereafter) was prepared with glutamine (Glutamax, Gibco 35050-038), antibiotics (Penicillin/Streptomycin, Gibco 15140-122), dexamethasone (200nM, Sigma, France) and 1% foetal bovine serum (Sigma F7524, lot BCBT8681). Culture flasks and permeable culture supports (Corning Transwell 3460, Corning, USA) were treated with collagen I (Sigma, dissolved in PBS, 100µg.ml^-1^, volume sufficient for 10µg.cm^-2^) by incubating the collagen solution on the surface overnight at 4°C. Excess collagen was aspirated prior to use and flasks washed with PBS. NCI-H441 and hAELVi cells were seeded at 4×10^4^ cells.cm^-2^ in collagen-treated flasks and cultivated in a humidified incubator at 37°C, 5% CO_2_.

For construction of models, Transwell® membranes (Corning 3460) were treated with collagen as described above, and cells were seeded at 1×10^5^ cells.cm^-2^. Other culture characteristics remained unchanged. Cells typically achieved full confluence within a week, as determined by thorough visual inspection, following which the apical compartment was drained and washed with HBSS with added calcium, magnesium and antibiotics (HBSS++) to remove serum proteins from the apical compartment. From this point on, models were cultivated at the air liquid interface, with medium supplied only in the basal compartment.

### Electrophysiological measurements

Transepithelial electrical resistance (TEER) was measured using an EVOM2 instrument (World Precision Instruments [WPI], USA) coupled with a cup electrode (ENDOHM-12G, WPI). Briefly, the electrode (lid and chamber) were sterilised with ethanol and left to dry aseptically in a laminar flow cabinet. The apical and basal compartments of the Transwell® insert were washed with HBSS++ to remove extracellular solutes and cell debris and 0.5mL of HBSS++ was added to the apical compartment. The electrode was filled with 2mL HBSS++, and connected to the EVOM2 unit. The insert was placed into the cup electrode with sterile tweezers and resistance measurements were collected at the first point when the value remained stable for 3 seconds. Resistance measurements from an empty insert cultured in parallel in the same media were subtracted from the measurements, and the values were normalised by multiplying by the surface area to give a resistance in Ω.cm^2^.

### Diffusion assays

Lucifer Yellow Carbohydrazide (Sigma L0259) was prepared as a 1mg.mL^-1^ stock solution in HBSS with calcium and magnesium (Gibco) and further diluted in HBSS to a working solution of 100µg.mL^-1^. Stale medium was aspirated, and both apical and basal compartments were washed with pre-warmed HBSS. 1mL of pre-warmed HBSS was added to the basal compartment of each well, while 0.5mL of LY working solution was added to the apical compartment. These plates were incubated in humidified conditions at 37°C, 5%CO_2_ for 1h to permit diffusion. A pairwise serial dilution of LY in HBSS from 100µg.mL^-1^ (50, 25, 12.5, 6.25, 3.25, 1.625, and 0.8125µg.mL^-1^) was prepared on a 96-well plate (black/clear, flat bottom, Greiner 655090) during the incubation period, along with triplicate samples of the original working solution and untreated HBSS. Samples of the basolateral medium at the end of the incubation were loaded alongside the standards and controls, and the plate was read on a Flexstation 3 (Molecular Devices) at 485nm_ex_/535nm_em_ with medium PMT gain.

The relationship between TEER and Lucifer Yellow diffusion was determined as described in Statistical Methods below.

### PCR

Primers were designed using NCBI primer-BLAST with the following changes to the default parameters: Product size 70-200bp, T_m_ 60°C<62°C<64°C, max T_m_ diff 2°C, primer pair separated by intron (1000-1000000bp), allow splice variants. Primers were accepted if positive controls generated a product in less than 40 cycles; matching the predicted size; with a single, well-defined T_m_ peak, and were redesigned if they failed to meet these criteria.

RNA was extracted from ALI cultures using NucleoSpin RNA mini kits (Machery-Nagel, Düren, Germany), RNA amount was measured using a nanodrop spectrophotometer (ThermoFisher Scientific) and reverse transcription was carried out using multiScribe High Capacity RT kits (Applied Biosystems, USA). All of these steps were carried out according to manufacturer’s instructions, and performed in immediate succession on ice to avoid RNA degradation. Samples of cDNA were stored at −20°C until needed for PCR. PCR was carried out in 384-well LightCycler 480 plates (Roche, Basel, Switzerland) in a LightCycler 480 thermocycler (Roche) using LightCycler SYBR Green I master mix (Roche). Primary analysis of the results was carried out in LightCycler 480 SW 1.5, and statistical treatment was carried out as described in the statistics section below.

### Western blotting

Protein was extracted in parallel to RNA extraction, using the Nucleospin kits (Machery-Nagel). Protein concentration was established using BCA kits (Pierce) and 10µg of protein was loaded into each well of a 4-20% pre-cast gel (BioRad, MiniProtean TGX), flanked with ladder at either end of the gel (PageRuler, ThermoFisher). Migrations were carried out in Tris-glycine buffer with 0.1% SDS at 100V for 1-2 hours as necessary, and transfer was carried out in tris-glycine + 20% methanol at 100v for 1h onto 0.45µm nitrocellulose membranes. Membranes were blocked for 1h with 5% milk in PBS-Tween, before primary antibody binding overnight at 4°C. Excess antibody was removed with multiple washes of blocking buffer, and HRP-conjugated secondary antibodies were bound for 30 minutes. After washing, bound HRP was visualised using ECL Prime visualisation reagent (Amersham, UK) on an AI600 gel imaging system (Amersham).

### Immunolabelling and confocal microscopy

Samples were washed with PBS and fixed by addition of 500µL of 4% PFA to the apical compartment for 10 minutes at room temperature (RT). For direct immunohistochemistry, the samples were washed with PBS, permeabilised by incubation with PBS +0.1% v/v Triton X-100 for 2 minutes, washed again with PBS and then blocked using 500µL PBS +2% w/v BSA for 1 hour at RT. Membranes were cut from the supports using a fine scalpel, placed face down on a 40µL drop of antibody solution (see Table 2) and incubated overnight at 4°C to permit thorough saturation of available epitopes. These samples were again washed and incubated with secondary antibody (Table 2) for 1 hour at RT. Samples were washed and mounted using Prolong Diamond (Invitrogen P36966), and imaged using a Zeiss LSM710 confocal microscope.

**Table 1:**
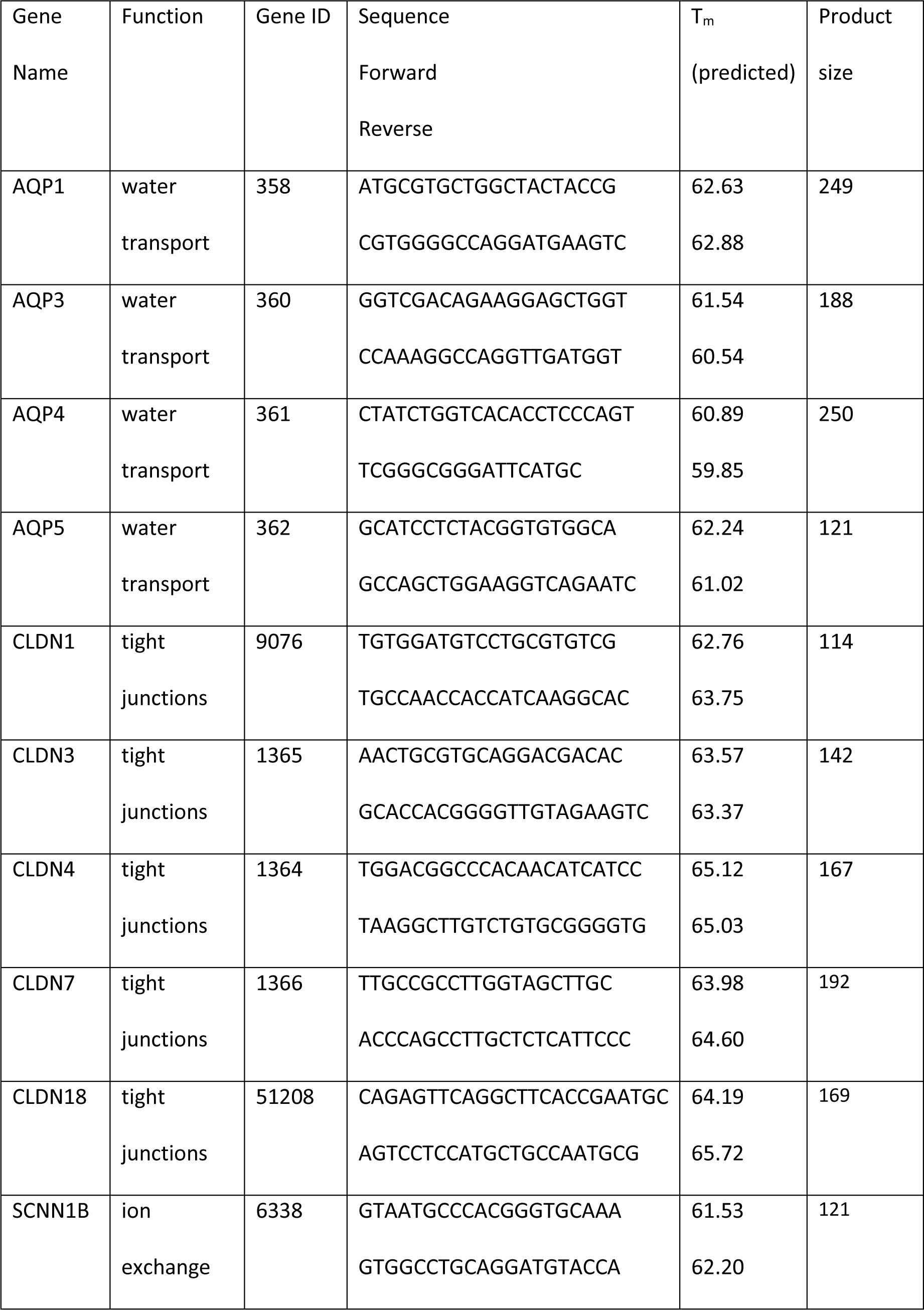
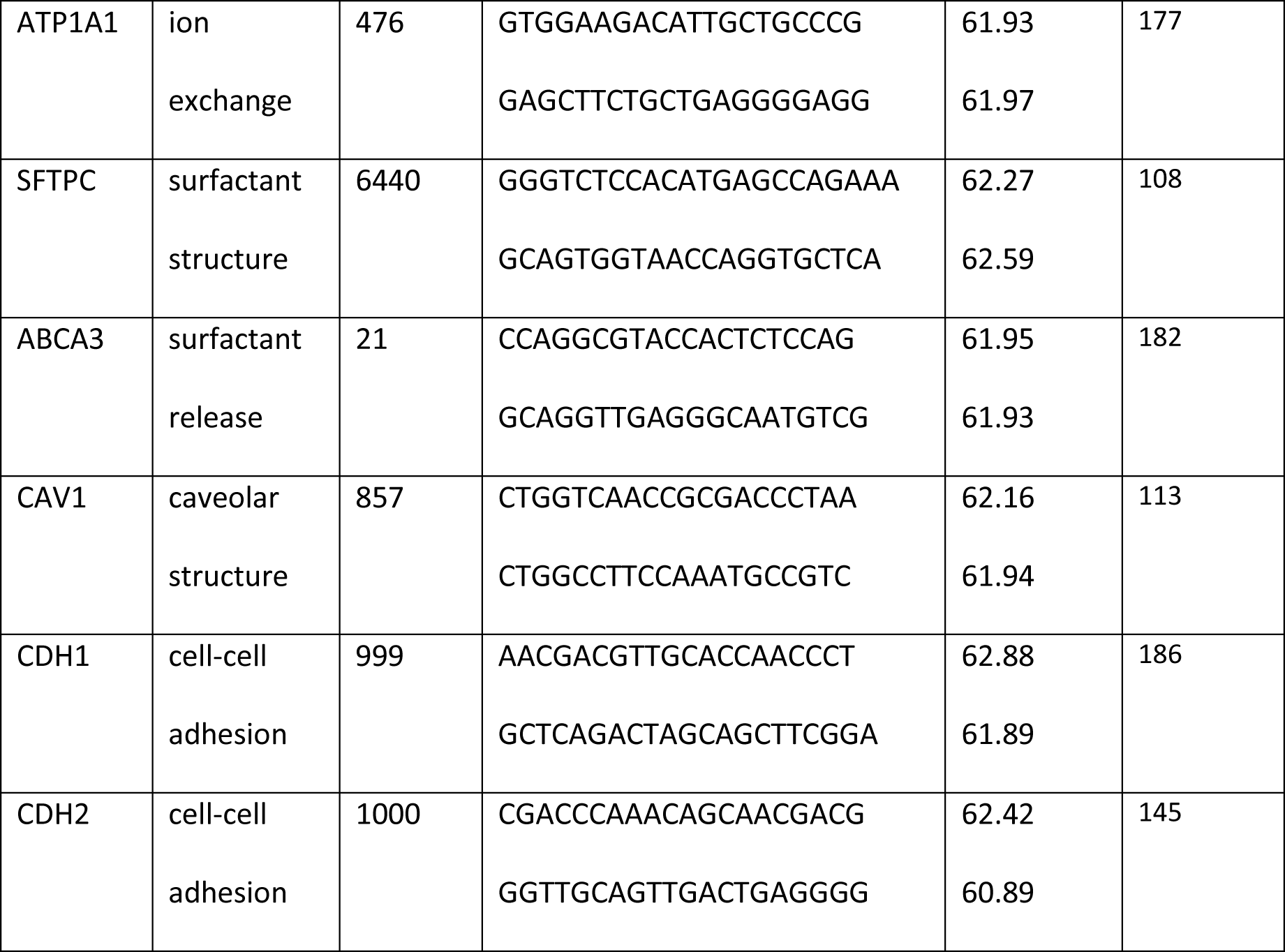
Primers used for experiments described in this paper.

**Table 2:** Antibodies used for experiments described in this paper.

| Target | Supplier | Ref/Clone name | Conc. | Secondary | Clonality |
| --- | --- | --- | --- | --- | --- |
| ZO-1 | Santa Cruz | H300 | 1:200 | DαR 488 | Mono |
| HTII-280 | Terrace Biotech | TB-27AHT2-280 | 1:50 | DαM 647 | Mono |
| Cav-1 | Sigma | SAB 4200216 | 1:200 | RbαM HRP | Mono |
| ABCA3 | ThermoFisher | PA5-52478 | 1:200 IF<br>1:1000 | DαR 488<br>RbαM HRP | Poly |
| Rabbit anti-mouse HRP | Jackson ImmunoResearch | 115-035-003 | 1:3000 | - | Poly |
| Donkey anti-mouse 647 | Abcam | ab150107 | 1:1000 | - | Poly |
| Donkey anti-rabbit 488 | Jackson ImmunoResearch | 711-545-152 | 1:1000 | - | Poly |

### Histological sectioning

After fixation in 4% PFA as described above, membranes were cut from their supports with a scalpel and dehydrated through a series of baths in increasing concentrations of ethanol in water (50%, 75%, 95%, 100%, 100%). After clearing 10 min in Clearene (Leica, Germany) to replace the ethanol, samples were immersed in liquid paraffin for one hour at 60 degrees to allow complete infusion of paraffin into the cells, and then set into block moulds and allowed to solidify at RT. Sectioning was performed on a Leica RM2135 microtome, and cut sections were deposited onto slides with the aid of a heated water bath. Prior to staining or immunolabelling, paraffin was removed with 2×10 min washes in Clearene, and then immediately rehydrated with the same series of ethanol baths in reverse order. For epitope retrieval, slides were immersed in citrate buffer at 90°C for 20min.

### Transmission electron microscopy

Barrier models, still mounted in the plastic Transwell® support, were fixed in 0.2M cacodylate buffer containing 2.5% glutaraldehyde and 2% paraformaldehyde for 45 minutes at RT. Each sample was postfixed, first in 1% OsO_4_ for 45 minutes at 4°C and then in 1% uranyl acetate for 2h at RT. Samples were briefly washed in water and then dehydrated through a series of increasing concentrations of ethanol (30%, 50%, 70%, 95% 100%) followed by 1:1 ethanol/propylene oxide and 100% propylene oxide, followed by embedding in Epon epoxy resin (Hexion). Ultrathin (80nm) sections were prepared using a Leica ultracut S microtome with a diamond knife (Leica Biosystems). Sections were contrast stained with lead citrate and photographed in a Jeol 100S transmission electron microscope (Jeol Europe) fitted with an Orius SC200 digital camera (Gatan-Roper Scientific).

## Statistical Methods

Statistical treatment was performed in Microsoft Excel and Graphpad Prism 8.

### Calculation of relative gene expression

Gene expression is calculated using the ΔΔCt method. Briefly, the Ct is calculated by the lightcycler software as the maximum second derivative of the fluorescence curve. The ΔCt value for each gene within each sample is calculated by taking the arithmetic mean of the differences between the Ct of the gene and the Ct of each of the reference genes for that sample (RPL13, RPL19 and HPRT). The ΔΔCt of a gene is calculated as the difference between the mean of the ΔCt of that gene for all replicates of that condition and the mean ΔCt of that gene for the hAELVi cell line replicates, effectively normalising the expression to hAELVi. Efficiency of each primer was measured empirically in preliminary experiments, and fold expression was calculated as 1 + efficiency raised to the power ΔΔCt.

### Mathematical description of the relationship between TEER and LY permeability

The reciprocal of TEER was taken as a starting point because electrical resistance is itself a reciprocal measurement of electrical conductance, where the charge carriers in this context are small soluble ions and the current is carried by diffusion of these ions. Additional terms were introduced to appropriately rescale the reciprocal graph to occupy the same range as the measured data (*a*), and to offset the minimum value (*b*). An additional power term was required to adjust the curvature of the model function to permit an acceptable R^2^ fit to observed data (*c*), which likely reflects differences in diffusion rate due to solute size. The model for the relationship between TEER and LY is thus as follows:

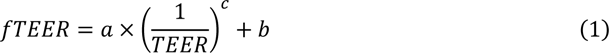

R^2^ values were calculated manually in Excel, and the Solver tool was used to adjust variables *a*, *b* and *c* to bring the R^2^ towards 1 (final values *a* = 26912.33, *b* = 681.92, *c* = 0.60, R^2^ = 0.98).

### Models of TEER development over time

TEER data were collected during routine medium changes throughout a 30-day period in ALI. These data were graphed to show TEER development over time. The reciprocal relationship between electrical resistance and molar permeability has the consequence that the variability of TEER measurements will increase in step with increasing TEER. Transforming the data logarithmically controls this heteroscedasticity, and there is uniform variance in logTEER. The graph of logTEER reveals a sigmoidal curve (as seen in **Erreur ! Source du renvoi introuvable.**), consistent with the logistic function which is commonly seen in biological growth. In order to fit the logistic model to the data, a logistic function was calculated in Excel over the range of 0-30 days using the general equation:

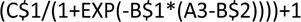

Where values A3-A34 contain the time in days, B1 will contain the doubling rate, B2 the midpoint of the curve (in days), and C1 the upper limit of log TEER. The +1 offset at the end of the expression equates to a minimum TEER of 10 once converted back to linear units, roughly equivalent to a transwell bearing non-barrier forming cells. R^2^ was calculated between this logistic function and logTEER, and the Excel Solver plugin was then used to optimise the parameters to provide the best fit (R^2^→1), shown in **Erreur ! Source du renvoi introuvable.**e and f as a black dotted line.

NCI-H441 exhibited an increase in TEER, followed by a decay. Reasoning that the same formative processes are likely responsible for TEER development, and that the decay was apparently irreversible, we reasoned that it may be possible to model the formation of “leaky patches” using a similar logistic function, antagonistic to the first. This is calculated as:

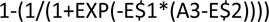

Where E1 and E2 will take the same roles as B1 and B2 in the previous expression, respectively. The function will reduce over time from 1 to approach 0. This is integrated with the primary logistic function by multiplying the two together with an offset variable. Again, Solver is used to optimise all the parameters to yield the best fit.

## Results

### Cell morphology and structure of epithelial cell lines

Phase contrast images were collected of hAELVi and NCI-H441 cells 6h and 4 days after seeding, in order to evaluate cell and colony morphology (**Erreur ! Source du renvoi introuvable.**). During exponential growth, both in flasks and on porous membranes, hAELVi cells demonstrate a compact individual morphology, extending cell processes but not spreading across the surface as might be expected for a cell type which is profoundly spread and flattened in vivo (**Erreur ! Source du renvoi introuvable.**a). NCI-H441 cells individually exhibit more uniform isotropic spreading, with fewer long processes (**Erreur ! Source du renvoi introuvable.**b). Quantification of the morphology of isolated cells in these images indicates that hAELVi cells have a larger mean spread area (**Erreur ! Source du renvoi introuvable.**c) and lower circularity (**Erreur ! Source du renvoi introuvable.**d). During subconfluent growth, hAELVi colonies are commonly found to have relatively loose association between the cells, many cells being partially or entirely isolated from nearby colonies (**Erreur ! Source du renvoi introuvable.**e). By contrast, NCI-H441 grow in compact clusters typical of epithelial cells with surrounding spaces largely devoid of individual cells (**Erreur ! Source du renvoi introuvable.**f). In an effort to quantify this, colony peripheries were manually identified in imageJ and measured. To control for varying perimeter:area ratios between colonies, this measure was normalised to colony cell number, the final result indicating that cells in hAELVi colonies exhibit a larger external perimeter per cell than those in NCI-H441 colonies (**Erreur ! Source du renvoi introuvable.**g), consistent with a looser colonial architecture and more convoluted cell shape.

Histological sectioning and light microscopy of hAELVi and NCI-H441 cultures indicates that, after 2 weeks of ALI culture, NCI-H441 cells amass to form a stratified layer of uneven thickness, commonly ranging from 30 to 50µm thick in a series of ridges and troughs across the area of a single transwell and a number of voids within the layer (**Erreur ! Source du renvoi introuvable.**a). By contrast, hAELVi cultures form thinner cell layers, typically around 10µm thick and consisting of a single cell layer (**Erreur ! Source du renvoi introuvable.**b), although stratification was observed in some replicates. These results were confirmed using electron microscopy (**Erreur ! Source du renvoi introuvable.** c and d show NCI-H441 and hAELVi, respectively). The cocultures typically appeared to form more compact layers than NCI-H441 single cultures of the same age, both macroscopically (personal observations, data not shown) and in thin sections viewed by electron microscopy (**Erreur ! Source du renvoi introuvable.**e). These results suggest that the cocultures are intermediate between the two parent lines in terms of the extent of stratification and layer depth.

TEM sections revealed the presence of inclusions or voids within the cell layers of ALI models, both in NCI-H441:hAELVi cocultures (**Erreur ! Source du renvoi introuvable.**a) and in NCI-H441 single cultures (Supplementary Fig S1). These voids were fully enclosed structures, not observed to connect with the apical surface or the underlying plastic membrane, and contained high concentrations of what appear to be irregular vesicles or cell debris. These voids were visible in 3D CLSM images of fixed ALI cultures, where immunolabelling of ZO1 revealed a pattern of staining consistent with tight junctions lining not only the apical surface, but also the interior of the intracellular voids (**Erreur ! Source du renvoi introuvable.**b). Similar subsurface voids were observed in FFPE sections of NCI-H441 ALI cultures in which the apical membrane of the cells lining the interior surface of the void feature HTII-280 (**Erreur ! Source du renvoi introuvable.**c), a large protein exclusively characteristic of the apical surface of alveolar type 2 epithelial cells in the native lung (9). Abundant projections closely resembling microvilli were observed on the interior surface of voids in both NCI-H441 and co-cultures (**Erreur ! Source du renvoi introuvable.**d). Furthermore, many small vesicles are present just below the lumenal membrane of these cells (arrowheads in **Erreur ! Source du renvoi introuvable.**d). These features collectively point to an apical identity for these domains of cells facing cavity lumina.

### Barrier function in epithelial monocultures and co-cultures

NCI-H441 and hAELVi exhibit distinct but robustly reproducible profiles of barrier development with respect to TEER. NCI-H441 cultures at ALI begin to develop a TEER which peaks at around day 10 and subsequently falls off (**Erreur ! Source du renvoi introuvable.**a). Models using hAELVi alone progressively strengthen their barrier over the first 15 days at ALI, after which development slows, but a high TEER is maintained until at least the end of the test period (**Erreur ! Source du renvoi introuvable.**b). Co-culture models, consisting of equal proportions of the two cell lines at the point of seeding, exhibit a pattern similar to that seen in hAELVi but with a slower development, taking some 21 days to develop a TEER comparable to the upper range of the hAELVi culture (**Erreur ! Source du renvoi introuvable.**c). In order to quantify this data more precisely, a mathematical model was fitted to the measured TEER data. The development of both hAELVi and coculture models, when plotted on a logarithmic scale, superficially resemble logistic growth, and indeed both are well described by a simple logistic function (R^2^ = 0.929 and 0.935, respectively). Postulating that the development of tight junctions in NCI-H441 might follow a similar fundamental logistic development which is progressively undermined by an irreversible degradation that spreads throughout the culture, the NCI-H441 logistic regression was multiplied by an inverse logistic curve ranging from 1 to 0. This model was fitted to the data, and the compound model was found to reasonably approximate the observed measurements (R^2^ = 0.719). From these models, it is possible to extract data concerning the limits and kinetics of barrier formation. The NCI-H441 cultures develop a maximum electrophysiological resistance of 137 ±5.8 Ω.cm^2^ which they reach approximately 7 days after establishment of air-liquid interface conditions, following which the resistance declines (**Erreur ! Source du renvoi introuvable.**d). The model indicates that the positive development of the barrier might reach 1238±52 Ω.cm^2^ in the absence of the degradation process. By contrast, hAELVi cultures approach a mean maximum resistance of 1439±62 Ω.cm^2^ (**Erreur ! Source du renvoi introuvable.**e). When grown in a mixed co-culture at the air-liquid interface, NCI-H441 and hAELVi interact to create a stable barrier with a TEER of 1758±74 Ω.cm^2^ (**Erreur ! Source du renvoi introuvable.**f), the highest observed value of all models tested. The mathematical models also suggest that formation of barriers in NCI-H441 is more rapid than in hAELVi (e, k=0.461 vs 0.335), but a slow decay process (k=0.224) attenuates this. Co-culture models exhibited a slower development than either parent line in isolation (k=0.249).

Consistent with the observed evolution of electrophysiological barriers, immunofluorescent labelling of ZO1 was observed to localise to tight junctions in both NCI-H441 and hAELVi (**Erreur ! Source du renvoi introuvable.**a). The permeability of tight junctions is controlled by the adhesion proteins of the claudin family, of which there is differential expression throughout the different cell types of the distal lung (10). A panel of several claudins (claudin 1, 3, 4 and 7) was quantified by RTqPCR in single and co-cultures at 14 days to characterise the nature of tight junctions in the two cell lines and identify any potential interaction between the two cell lines with respect to junction composition in the co-culture (**Erreur ! Source du renvoi introuvable.**b). Claudin 1 transcripts were present at detectable levels in all cell lines, the coculture expressing claudin 1 at a level that is not significantly different from that of hAELVi, while expression was slightly higher in NCI-H441. Claudin 3 is expressed ∼10-fold more strongly in NCI-H441 cells than in hAELVi, which is consistent with observations of claudin expression in native type II cells compared to their type I counterparts. Claudin 3 expression in cocultures at day 14 appears to be intermediate between the two parent lines. Claudin 4 and 7 exhibit similar patterns of expression, the coculture and NCI-H441 being statistically similar, whereas hAELVi expresses these genes at a lower level. Another important mechanism which supports pulmonary function is the clearance of water from the distal airways. This occurs by movement of water through aquaporin channels to balance osmotic pressure established by active transport of solutes out of the pulmonary fluid (11). To investigate the expression of genes related to water homeostasis, RTqPCR was carried out after 14 days of culture using primers for aquaporins known to be expressed in the lung (AQP1, AQP3, AQP4 and AQP5). Among these, aquaporin 4 is distinctly differentially regulated, being around 3 orders of magnitude more strongly expressed in NCI-H441 than in hAELVi cultures, while co-cultures exhibit intermediate transcript abundance. In addition, aquaporins 1 and 5 are also more abundantly transcribed in NCI-H441 than in hAELVi cultures, co-cultures exhibiting levels of expression similar to NCI-H441. In contrast, AQP 3 is expressed at higher abundance in hAELVi cultures (**Erreur ! Source du renvoi introuvable.**c) than in NCI-H441, and co-cultures exhibit an intermediate level of expression. Finally, the genetic basis for the ion exchange which drives fluid clearance from the distal airspaces was interrogated, using primers of the two most important ion channels for fluid homeostasis in the lung, ENaCB1 and ATP1A1. These two genes are both expressed more strongly in NCI-H441 relative to hAELVi cells. ATP1A1 expression in the coculture is similar to that of hAELVi, while ENaCB1 expression in the co-culture is intermediate between that of the two parent lines.

### Surfactant production in NCI-H441 monocultures and co-cultures

FFPE sections of normal human lung tissue show that type-II cells jointly express the apical type-II antigen HTII-280 and ABCA3, a component of the lamellar body membranes responsible for surfactant release (**Erreur ! Source du renvoi introuvable.**a). ABCA3 is also observed within the cells of NCI-H441 ALI cultures (**Erreur ! Source du renvoi introuvable.**b), supporting the use of the line as a model of the surfactant-secretory phenotype. HTII-280 is also observed at the apical surface of the cell layer (**Erreur ! Source du renvoi introuvable.**c), further supporting the inference of type-II identity of NCI-H441. TEM observations of ultra-thin sections of ALI cultures also revealed the presence of lamellar bodies in NCI-H441 (**Erreur ! Source du renvoi introuvable.**d), which were not observed in hAELVi or cocultures. Expression analysis of NCI-H441, hAELVi and cocultures after 14 days of culture in ALI conditions by qPCR revealed a profound difference between hAELVi and NCI-H441 in terms of surfactant-associated protein C (SFTPC) gene expression, with NCI-H441 cells exhibiting SFTPC transcripts at a level several orders of magnitude higher than hAELVi cells. Analysis of ABCA3 gene expression showed also a ten-fold greater transcript abundance in NCI-H441 than hAELVi cultures (**Erreur ! Source du renvoi introuvable.**e). These observations are consistent with the proposed identity of the two cell lines. Notably, the expression of both transcripts in the coculture is intermediate between the two parent cell lines, implying the continued co-existence of the two cell lines in the model over at least two weeks of culture. Western blotting using a polyclonal antibody against ABCA3 (**Erreur ! Source du renvoi introuvable.**f) revealed multiple bands which are consistent with the proteolytic cleavage reported by others, which is reported to be linked to ACA3 trafficking and potentiation (12). The full-length protein, expected size 190kDa, and the large cleaved fragment, expected at 150kDa, both appear larger on our gels compared to the PageRuler ladder that was used for reference (250kDa and 180kDa, respectively). As this was seen in both cell lines, it is more likely an artefact than a real deviance of the ABCA3 protein in these cells from previously reported sizes. The 40 kDa band is likely the small cleaved fragment whereas the identity of the 50kDa band is unknown. Interestingly, the 50 kDa fragment is differently sized in the two parent lines, and bands of each of these variants are seen in the lysate of 14-day cocultured ALI models, again suggesting that the coculture stably persists for this duration.

### Caveolae in hAELVi and coculture are post-transcriptionally regulated

Caveolae are observed by TEM (**Erreur ! Source du renvoi introuvable.**a), although these structures were only readily identified in a minority of cells in the coculture. We anticipated that caveolin-1, which is an important structural component of caveolae, would be expressed more strongly in hAELVi, consistent with published results for type I vs type II cells (13). However, although we observed little difference in expression of caveolin-1 between the two cell lines or the coculture (**Erreur ! Source du renvoi introuvable.**b), WB analysis showed that caveolin-1 protein was more abundant in hAELVi, with NCI-H441 and co-culture models exhibiting much lower protein abundance after 2 weeks of culture at ALI (**Erreur ! Source du renvoi introuvable.**c and d).

## Discussion

There is a clear need for models of the alveolar barrier to support ongoing research in toxicology and therapeutics. It is important that future models reflect the diversity of the tissue in order to surpass the limitations of single-cell type models. The two alveolar epithelial phenotypes make up one aspect of this diversity. Type I cells are profoundly flattened, and therefore cover 95% of the alveolar surface, despite only constituting half as many cells as the columnar type II cells (14). There is an ontogenic relationship between the two, in that Type II epithelial cells terminally differentiate into Type I cells (15), although recent research indicates that both type I and type II cells comprise of multiple subtypes with differing degrees of developmental plasticity (16–19), with their own niches and supporting cell types (20–22). Future research is likely to reveal further complexity within the epithelial landscape, and models will have to evolve to reflect this knowledge if they are to be truly representative of the native tissue. In this paper we have aimed to not only further characterise two cell lines with respect to their individual properties in an ALI model system, but to investigate the potential interaction between these cell lines with a view to incorporating this diversity in future models of the barrier.

The cell lines that were selected for this study were chosen on the basis of their reported phenotypic similarity to type I and type II cells. The cell line A549, which is perhaps the most widely used cell line in models of the distal lung, was not selected because it is reported to be a poor representative of type II cells in terms of phenotype-specific gene expression (23, 24) and was found in preliminary tests not to develop a TEER that was significantly greater than non-barrier cell lines. Instead, we selected NCI-H441, which has been reported to express surfactant proteins (25, 26) and to form a better electrophysiological barrier than A549 (26, 27). It is more difficult to find representative models of type I alveolar epithelial cells. Primary type I cells exhibit limited self-renewal in culture, and there is not an abundance of type I-derived cancer cell lines available from commercial repositories. However, an artificially immortalised cell line, hAELVi, was recently created from primary type I cells to address this shortfall (8). This cell line forms a good paracellular barrier and exhibits gene expression that is broadly comparable with the parent cell type (8).

The two cell lines use different media, so it was necessary for coculture to bring the two lines into a common medium. We found in preliminary tests that hAELVi performed poorly in air-liquid interface culture using simple media such as RPMI 1640, in which NCI-H441 is typically grown, but that both cell lines attained reasonable TEER values in advanced DMEM (aDMEM), although cell adhesion and proliferation was dependent on the addition of a small amount of serum (1% FBS). As we required a medium with which both cell lines were compatible, we therefore selected this as a working medium. It was also necessary to add a glucocorticoid receptor agonist (dexamethasone or hydrocortisone) to stimulate barrier formation, as reported previously by others (26).

One attribute which differs between these cells in the culture models, individually or in conjunction, and alveolar cells in vivo is cell morphology. ATI cells in the human lung individually cover an area of the alveolar surface of about 5,000 µm^2^ and have an average thickness of less than one micron (14), while cells in the culture model, even in the absence of stratification, are considerably less spread and deeper in section. This is not due to physical crowding as a result of the hyperproliferation, as subconfluent cells do not exhibit this profoundly flattened and spread morphology prior to the onset of crowding. Cell spreading often occurs in other contexts as a result of applied biomechanical forces, or as a response to the mechanical rigidity of the substrate. The cell lines we use in our model have been shown to exhibit other physiological responses to stretching forces (28), and the mechanical properties of the plastic membrane are very different from the extremely deformable environment of the alveolar wall. The composition of the underlying matrix may also influence the development of morphology, and these cell lines may spread better on a surface that includes specific components of the basement membrane such as collagen IV (29), collagen VI (30) or Laminin-311 (31). Interactions with other cell types may also drive morphological development of the epithelial cells, not least of all by depositing different matrix proteins or influencing the composition of the matrix deposited by epithelial cell cultures (32). Cell morphology is an accessible, though indirect, indicator of cellular phenotype and we have presented data in this paper to establish a baseline for future models. Extending the complexity of the coculture to incorporate other relevant cell types and contextually appropriate biomechanical stimuli may help to bring the morphology of cells in *in vitro* models like ours towards that seen in human lung epithelium *in situ*.

Although most of the alveolar epithelium is a simple squamous epithelium in vivo, stratification is routinely observed in NCI-H441 and is seen rarely in hAELVi cultures, probably due to decreased contact inhibition in the transformed cell line. However, we have noted the existence of some unusual cavities within stratified cell layers in NCI-H441 and mixed coculture models indicating that this cell line possesses intrinsic mechanisms which act to maintain polarity even when fully enclosed by other cells. Indeed, we presented several pieces of evidence that these cavities represent an additional apical surface with respect to the polarity of the surrounding cells. These internal surfaces bear microvilli, are sealed with tight junctions and possess membrane-bound vesicles at the surface, suggesting that they actively transfer material into and/or out of the cavity lumen. The apical character of the lumen-facing cellular domains imply cell polarity, suggesting that there may be a corresponding additional basolateral domain surrounding these features, at regions of contact with the true apical cells of the model. The persistence of this polarity-forming mechanism further supports the use of NCI-H441 as a model line. This also suggests that cocultures of hAELVi and NCI-H441 might be amenable to forming organoids in suspension culture, where the intrinsic pattern-forming behaviour is not constrained by artificially imposed conditions such as a rigid synthetic substrate. However, the presence of these structures also indicates that cell polarity is not restricted to the plane of the culture surface in cultures containing these features, and therefore active transport of water, solutes and suspended nanomaterials across ALI models cannot be assumed to be uniformly directed between the macroscopic apical and basal compartments. This therefore represents a potential confounding factor in historical studies using NCI-H441 in permeable supports, and a way to prevent stratification and cavity formation would increase the power of these models to represent polarity-dependent characteristics of the alveolar epithelium.

There is a substantial difference between the two alveolar epithelial lines in the long-term development of TEER. In contrast to the progressive development of TEER exhibited by hAELVi seen in our own results and those of others (8), NCI-H441 TEER development in our experiments reaches a peak and then subsides to a much lower resting value, similar to trends seen in existing literature concerning the cell line (26, 27). To analyse this biphasic response, we created a simple mathematical model of TEER to evaluate the kinetics of this process. Modelling positive TEER development in NCI-H441 with a logistic function, as observed in hAELVi, an additional component is needed to account for the subsequent negative change in TEER, which can be described with a negative logistic function. In this case, the best-fit values suggest an initial formation of tight junctions comparable to hAELVi, which is undermined by a slower, progressive decline. This could be related simply to proliferation and overcrowding of the cancer-derived line, or might be the result of changes within the tight junction such as progressive incorporation of pore-forming claudins over time (33). It is also possible that changes in barrier permeability occur as a result of signalling independent of changes in gene expression, as has been observed in primary rat lung epithelium (34), perhaps as a result of phenotypic changes during ALI culture. It is noteworthy that co-cultures also go on to develop a TEER comparable to that of the hAELVi monocultures and which do not decay in the timescale over which we observed them. Taken independently, these observations could suggest that hAELVi cells come to dominate the culture either in terms of population or in terms of their influence on the formation of the barrier, but gene expression indicative of NCI-H441 cells persists as the co-culture develops. The continued presence of both NCI-H441 and hAELVi in the co-culture may hint at some reciprocal regulatory interaction between the two lines with respect to proliferation and quiescence as is reported to exist in mice lung development (35).. The elevated TEER in presence of NCI-H441 leads us to speculate that, while NCI-H441 cells do not form particularly strong barriers between themselves, they may form stronger and more stable tight junctions with neighbouring hAELVi, an interaction that would be commensurate with the distribution of type II cells in the lung, where they are interspersed between type I cells.

As the permeability of tight junctions in the lung and elsewhere is largely controlled by claudins (10, 36), we investigated their expression in support of our electrophysiological characterisation. The most abundant claudins in the distal lung are understood to be claudins 3, 4, and 18 (34). Claudin 1 is not anticipated in the alveolar epithelium in significant quantity but is abundant in lung tissue as a whole (10), and therefore may reflect de-differentiation, perhaps as an artefact of extended in vitro culture or a result of transformation and immortalisation. We observed only minor differences between the two cell lines in terms of their expression of claudins 1, 4 and 7, but it is interesting that the coculture is simultaneously more similar to hAELVi in terms of claudin 1 expression, and closer to NCI-H441 in claudin 4/7 expression. This suggests a level of differential regulation of one or both genes, without which we would expect all genes to be expressed in the coculture at a level intermediate between the two parent lines, dependent only on the ratio of the two cell types in the coculture at the point of RNA collection. Interestingly, claudin 7 is understood to have significant impact in cancer pathology and cell migration (37, 38), and we anticipated that it may therefore be downregulated in NCI-H441, which is derived from a small-cell adenocarcinoma. Interestingly we observed slightly higher claudin 7 expression in NCI-H441 compared to hAELVi cultures, and understanding this anomalous finding may help to shed further light on the role of claudin 7 in pulmonary adenocarcinoma. Claudin 3 expression is the most divergent between the two parent lines, NCI-H441 expressing claudin 3 at a ten-fold higher level than hAELVi, consistent with their respective identities as type I and type II AECs (34).

We also observed a profound difference in expression of genes associated with surfactant production (SFTPC, ABCA3), both genes being expressed at a substantially higher level in NCI-H441 than in hAELVi, which is consistent with the perception of the former cell line as representative of type II pulmonary epithelial cells while the latter is representative of type I. The high level of expression of SFTPC in the coculture again underlines the persistence of the type II cell line within this culture model. Furthermore, the presence of lamellar bodies in the cytoplasm of NCI-H441 cells confirms that surfactant is actively being produced and transported.

Water homeostasis in the alveolar space is driven by active transport of solutes, especially small ions, across the epithelium from the alveolar lumen into the blood stream. This establishes an osmotic pressure leading to net diffusion of water out of the alveoli (reviewed in (39)). Solute transport is mediated by a suite of different transporters, among which the sodium transporter ENaC1 and the potassium transporter ATP1A1 are particularly important (39–41). The water itself moves via aquaporin channels, several of which are selectively expressed in different regions of the lung (42). Expression analysis covered a panel of lung-specific aquaporins known to be differentially expressed throughout the lung (AQP1, AQP3, AQP4 and AQP5) (42). AQP1 expression is normally characteristic of endothelial cells, and its presence here is therefore unexpected. It has been reported that expression of AQP1 is absent in healthy alveolar epithelium, but is observed in biopsies from patients with idiopathic pulmonary fibrosis (43). AQP1 was also observed to be induced by TGFβ signalling in A459 cells, suggesting a mechanistic element that could be investigated by disrupting TGFβ signal transduction. AQP3 expression is reported in type II cells and absent from Type I, and it is therefore surprising to see higher expression in hAELVi than in NCI-H441. Aberrant patterns of aquaporin expression may be a feature of one of the cell lines, or might arise from discrepancies between the native tissue environment and the culture conditions. Similar unexpected patterns of expression are seen for AQP4 and AQP5. AQP 4 expression is highly divergent, as expected, but is abundant in NCI-H441 cells whilst being substantially less highly expressed in hAELVi. Again, this is contrary to expectations and raises questions about phenotypic identity. While AQP5 is significantly upregulated in type I epithelial cells, and aquaporin 5 therefore probably plays a significant role in water clearance in vivo, it has already been noted that expression of AQP5 by hAELVi is surprisingly weak (8), and our results support this. The expression of AQP5 has also been noted in ATII cells (11), and it is therefore not surprising that AQP5 expression might be higher in NCI-H441 than in the AQP5-deficient hAELVi cells. In the cocultures, intermediate expression of aquaporins 1,3 and 4 were observed, relative to the parent lines. Again, this is generally consistent with the persistent presence of both cell lines in the milieu. Only AQP5 exhibits a different signalling pattern, RNA abundance being indistinguishable from that of NCI-H441 alone. If we accept the hypothesis that both cell lines persist to at least this stage, this indicates that AQP5 expression is upregulated in one or both parent lines as a result of interactions in the coculture.

Caveoli are an ultrastructural feature of the apical membrane which are reportedly present in type I epithelial cells and absent from type II cells (44). Furthermore, expression of the CAV1 gene encoding caveolin 1, the principal structural protein and driver of caveolin formation, is reported to be absent from primary type II cells and upregulated during spontaneous differentiation in culture (13). In our experiments, we were able to observe caveoli in co-cultures but not in single cultures of NCI-H441 or hAELVi. Surprisingly, we found that expression of the CAV1 gene did not vary significantly between the two parent lines. However, western blotting revealed a marked difference in abundance of caveolin 1 protein in hAELVi, suggesting that there is some level of post-transcriptional control of expression of this protein. The co-culture appears to express CAV1 protein at a low level similar to NCI-H441 single cultures. The presence or absence of caveolin 1 in alveolar epithelial models may have diverse consequences, as the full role of caveoli in the native lung is not yet fully understood. Recent research has found that caveolin-specific proteins are indispensable for mechanotransduction of mechanical stretch in type I cells, and that the induced response includes calcium-mediated paracrine signalling which stimulates lamellar body exocytosis in type II cells during co-culture (28). The presence of caveoli may therefore be essential in future models which include biomechanical stimuli as well as coculture of cells of multiple lineages. Caveolin-1 has also been linked to endocytosis and albumin transport in other contexts (45, 46), which may have ramifications on susceptibility of the model to toxic compounds or nanomaterials.

In conclusion, we have established a cell culture model of the alveolar epithelium composed of commercially available cell lines and materials which features co-culture of cells representative of type I and type II pneumocytes which are maintained over 21 days making this co-culture system a very promising alternative model to study the effects of toxic compounds and therapeutics. We have established baseline values for TEER and mRNA expression of a panel of phenotypically important genes that will facilitate experimental design by other researchers wishing to use this model. We have also identified intracellular voids that form in NCI-H441 cultures over long-duration culture at the air-liquid interface, which may have relevance to interpretation of past or future results of experiments using this cell line. These features demonstrate apical basal-polarity formation, and may therefore present unique research opportunities. We also noted the presence of ultrastructural features in the coculture model which are unambiguously linked to phenotypic identity of the cells, such as lamellar bodies and caveoli. Finally, we observed evidence of post-transcriptional regulation of caveolin-1, implying that this model may be a suitable platform for investigating this potentially novel mechanism of caveolin regulation.

## Supporting information

20210802_SI-BioRxiv-OB

## Acknowledgements

This research was supported by the Agence Nationale de la Recherche under the contract ANR-17-CE09-0017 (AlveolusMimics). The authors would like to express our gratitude to Jamileh Movassat for allowing us the use of her laboratory and equipment for paraffin embedding and sectioning of samples. We also express our gratitude to Imagoseine and to the BFA imaging platform, for the use of their confocal microscopes. Finally, we would also like to express our gratitude to Pascal Roussel and Valentina Sirri-Roussel for their assistance with western blotting.

---

**Fig 1.**
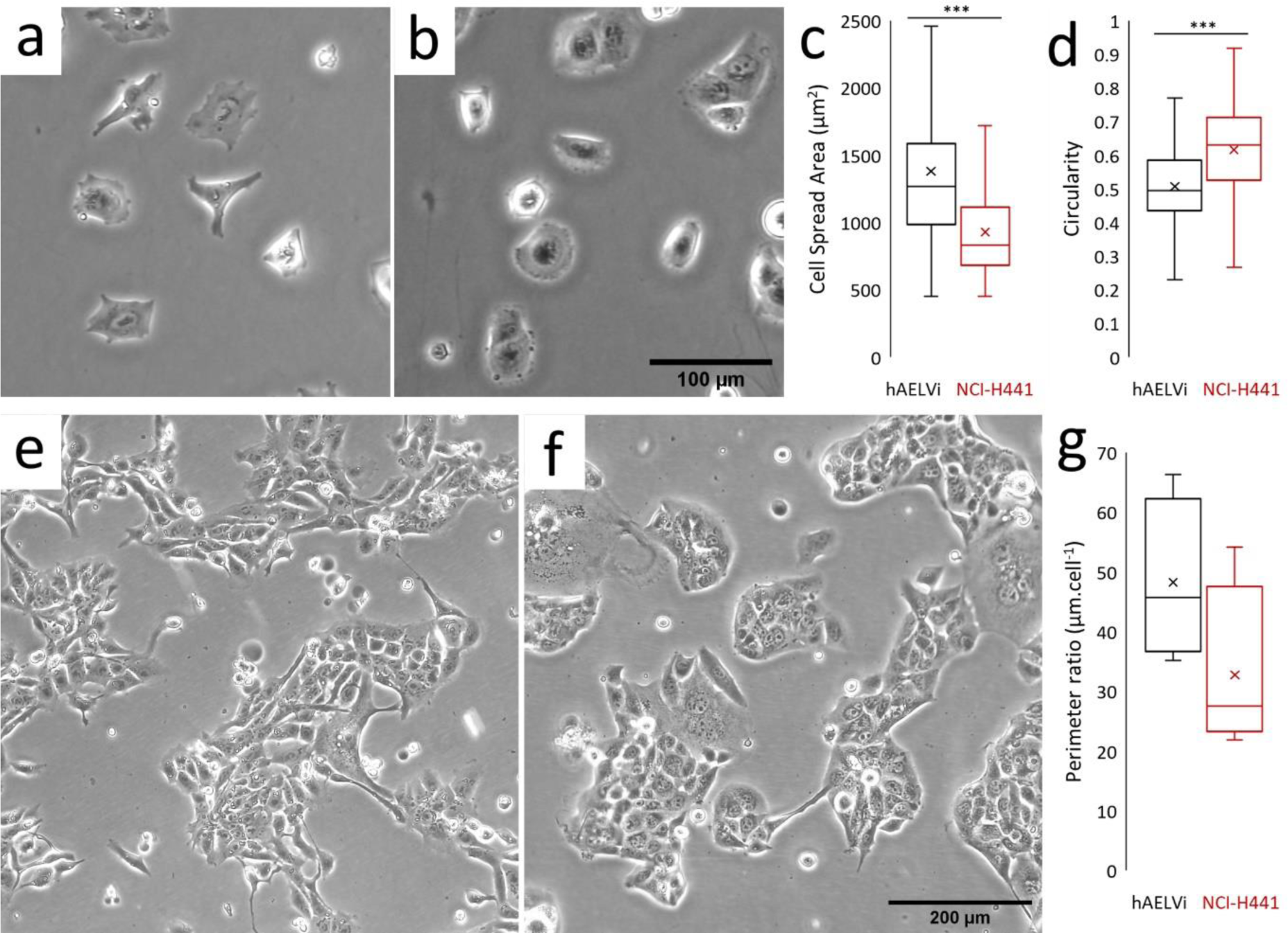
Comparison of the morphology and growth habit of hAELVi and NCI-H441. Phase contrast images of submerged cultures of hAELVi (a, e) and NCI-H441 (b, f) cells, at time points 6 hours (a, b) and 4 days after seeding on plastic culture flasks (e, f). Single cells were analysed within the captured images to determine cell spread area (c) and circularity (d) using ImageJ. Colony sizes were determined by measuring the length of the perimeters of colonies and normalising this value to the number of cells in the colony (g). *** = P<0.001, n=15 or more cells in 5 or more randomly selected fields.

**Fig 2.**
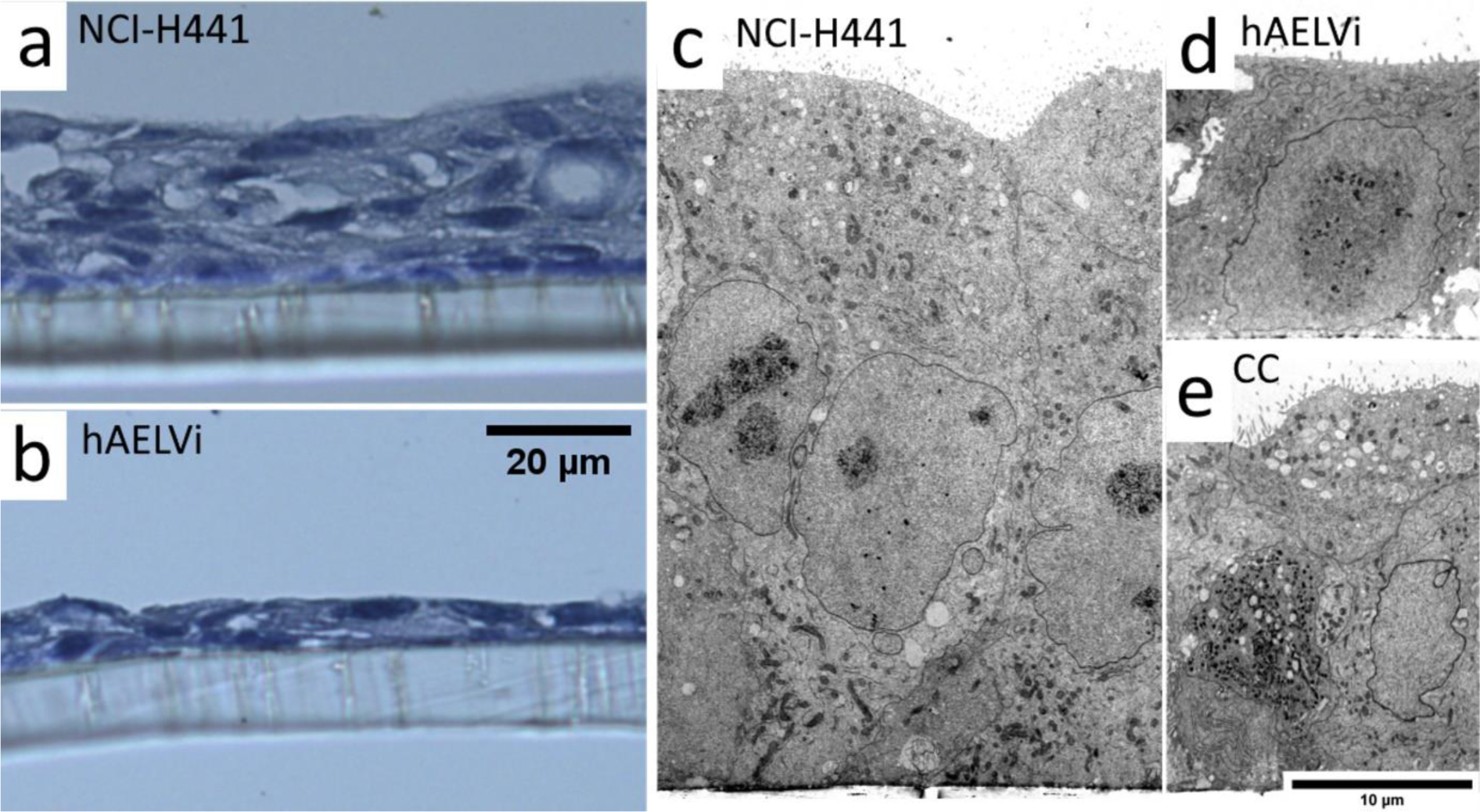
Supra-cellular structure of ALI cultures. Light microscopy images of ultrathin sections stained with Papanicolaou stain (a, b) and TEM images (c, d, e) of ALI cultures of NCI-H441 (a, c) or hAELVi (b, d) or the coculture (e) after 14 days of culture. NCI-H441 cultures exhibit a loose stratified cell layer, while hAELVi typically form a flatter monolayer. Cocultures do exhibit stratification, but are generally more consistent than NCI-H441 alone.

**Fig 3.**
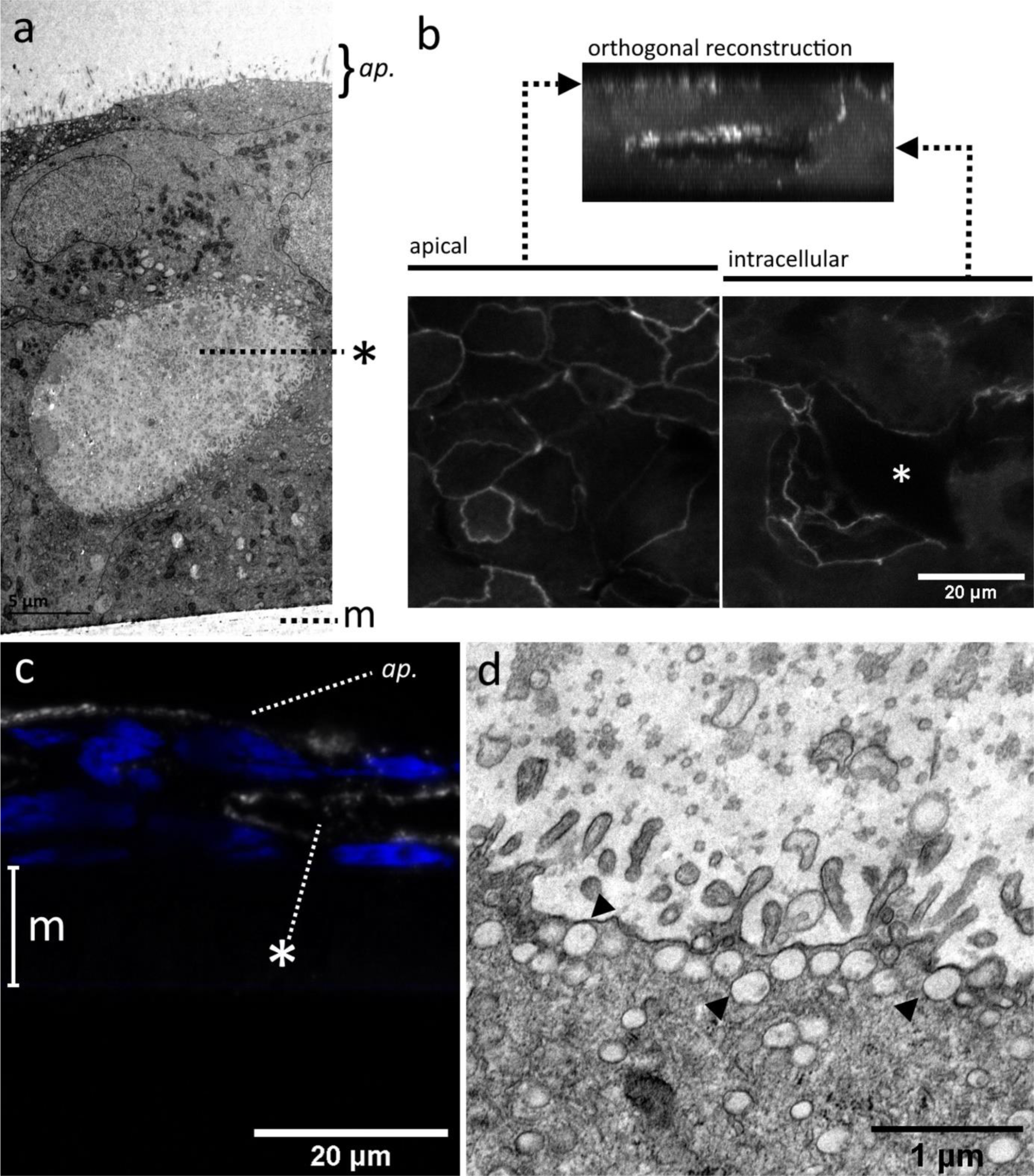
TEM images of voids within stratified epithelial layers. TEM images of NCI-H441:hAELVi cocultures grown at the air-liquid interface for 2 weeks reveal the existence of cavities beneath the surface (a). CLSM images of ZO-1 immunolabelled samples (b) indicate that these structures are lined with tight junctions on their luminal surface, in addition to those sealing the apical surface. Similar structures seen in FFPE sections of NCI-H441 grown in ALI conditions stain positively using an antibody against HTII-280, an antigen found exclusively on the apical membrane of type II epithelial cells (c). Voids in NCI-H441 single cultures and cocultures are lined with microvillous processes and secretory vesicles, also features of the apical surface of type II alveolar epithelium (d). Asterisks mark internal voids. Polymer support membrane labelled m where visible. ap. = apical surface.

**Fig 4.**
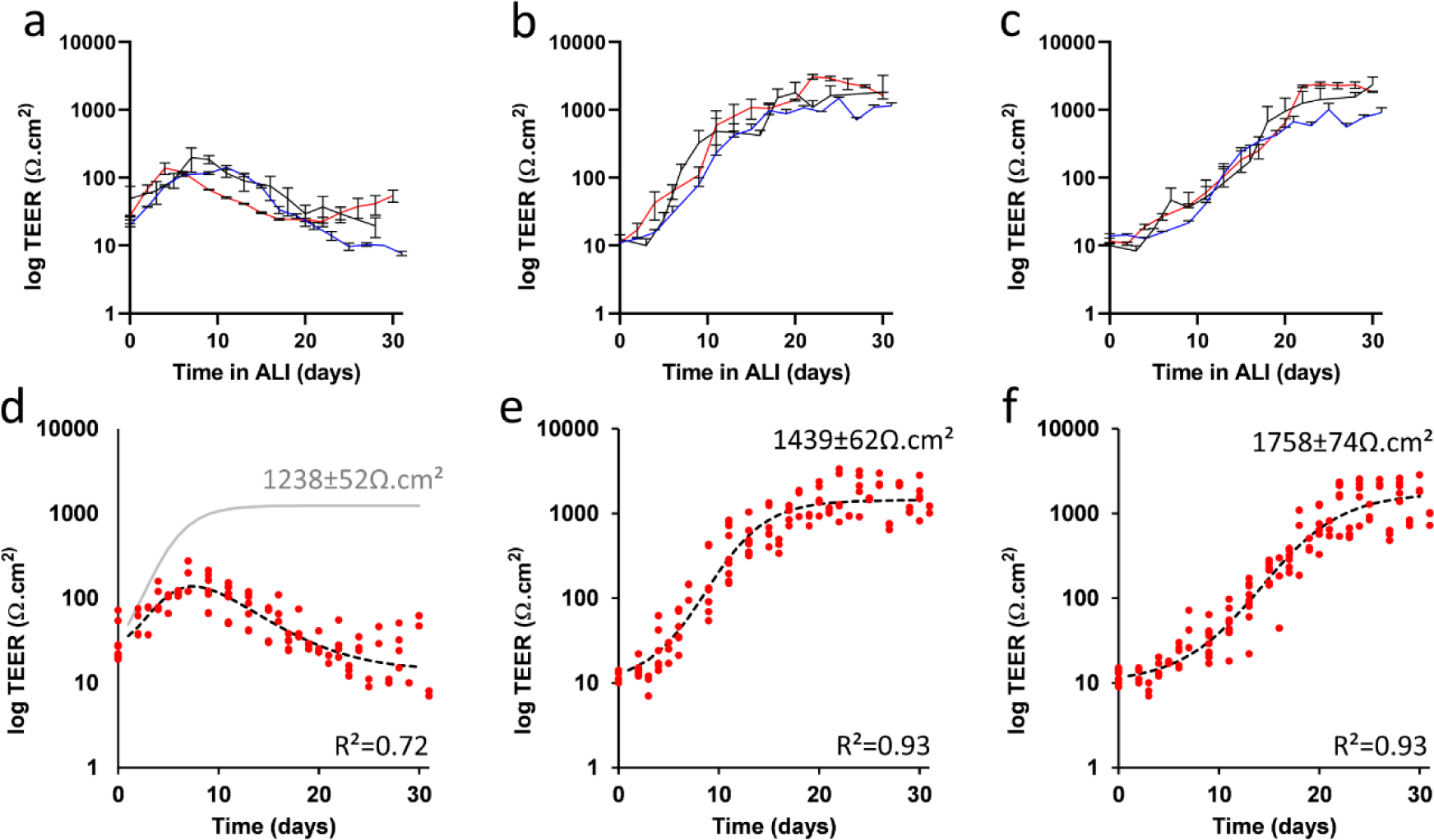
Electrophysiological barrier formation of NCI-H441, hAELVi, and co-culture in ALI conditions. TEER measurements of NCI-H441 (a), hAELVi (b) and an equal co-culture of the two (c) over a period of thirty days following the establishment of ALI conditions. (3 independent experiments, n=3 transwells). These data were fitted by regression to a logistic growth and decay model (d,e,f, red points indicate data values summarised in a,b and c, respectively). Only the model for NCI-H441 (d) includes a non-zero decay term; the dotted line shows the best fit including this decay term, while the grey line shows the logistic growth component in isolation for comparison to hAELVi (e) and co-culture (f). In preliminary experiments, we observed a direct mathematical relationship between TEER and Lucifer Yellow (LY) diffusion (Supplementary Fig S2), suggesting that TEER and actual diffusion assays are mutually redundant, at least for LY, and that no new information is gained by performing both measurements.

**Fig 5:**
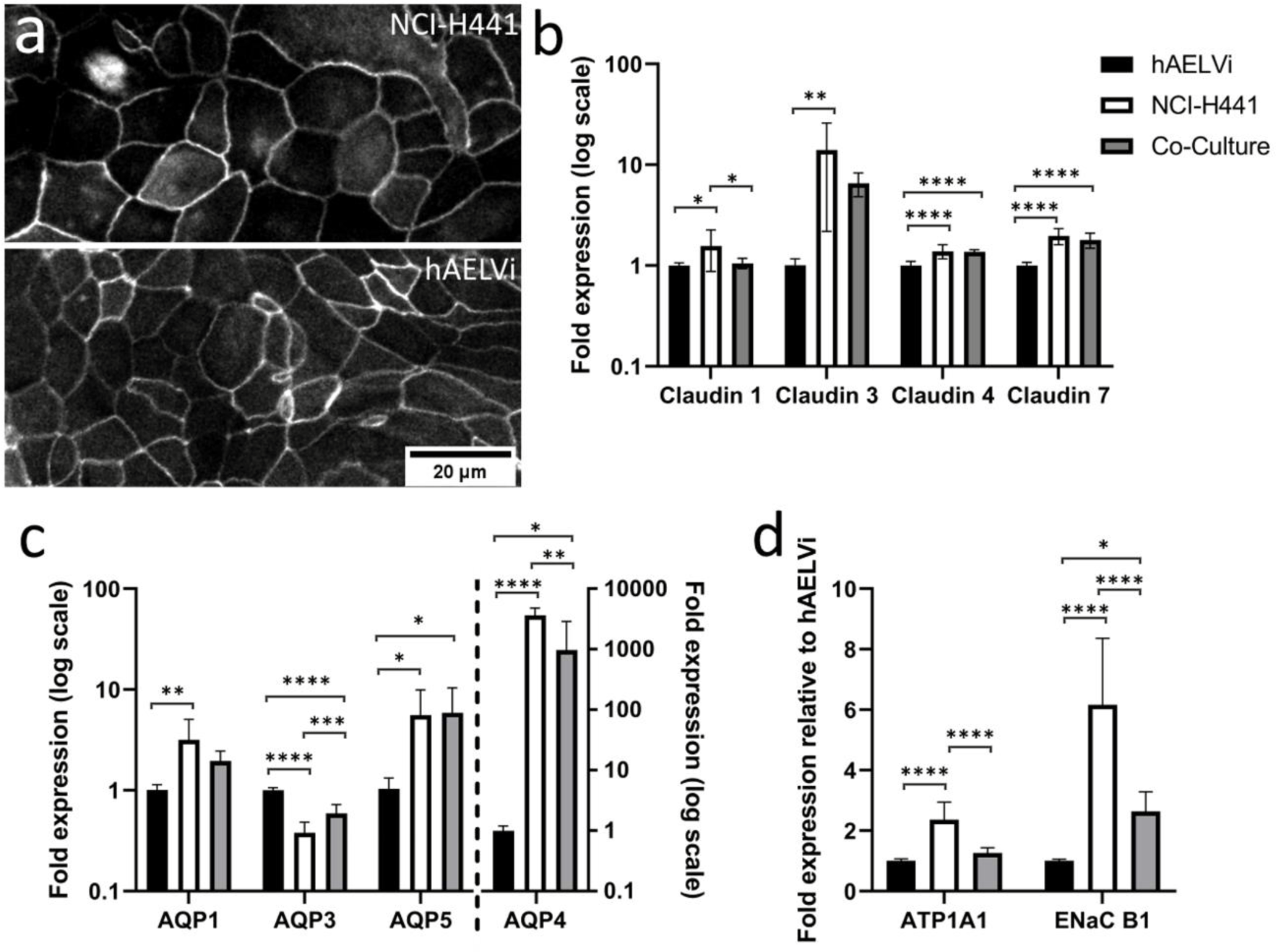
Differences in expression of key functional genes between hAELVi, NCI-H441 and co-culture. Cultures of hAELVi, NCI-H441 cells and a co-culture consisting of equal parts of both cell lines were maintained in ALI conditions for two weeks after confluence. Immunolabelling of ZO1 in the two parent lines revealed localisation to the cell periphery, consistent with incorporation into tight junctions (a). Gene expression analysis by RTqPCR of tight junction proteins (claudin 1, 2, 3 and 4; b), water channels (aquaporins AQP1, AQP3, AQP5 and AQP4 (c)) and ion pumps (ATP1A1 and ENaC B1; d) in hAELVi, NCI-H441 and a co-culture (fold expression relative to expression in hAELVi after normalisation to reference genes RPL13, RPL19 and HPRT. n=3 replicates within 3 independent experiments.

**Fig 6:**
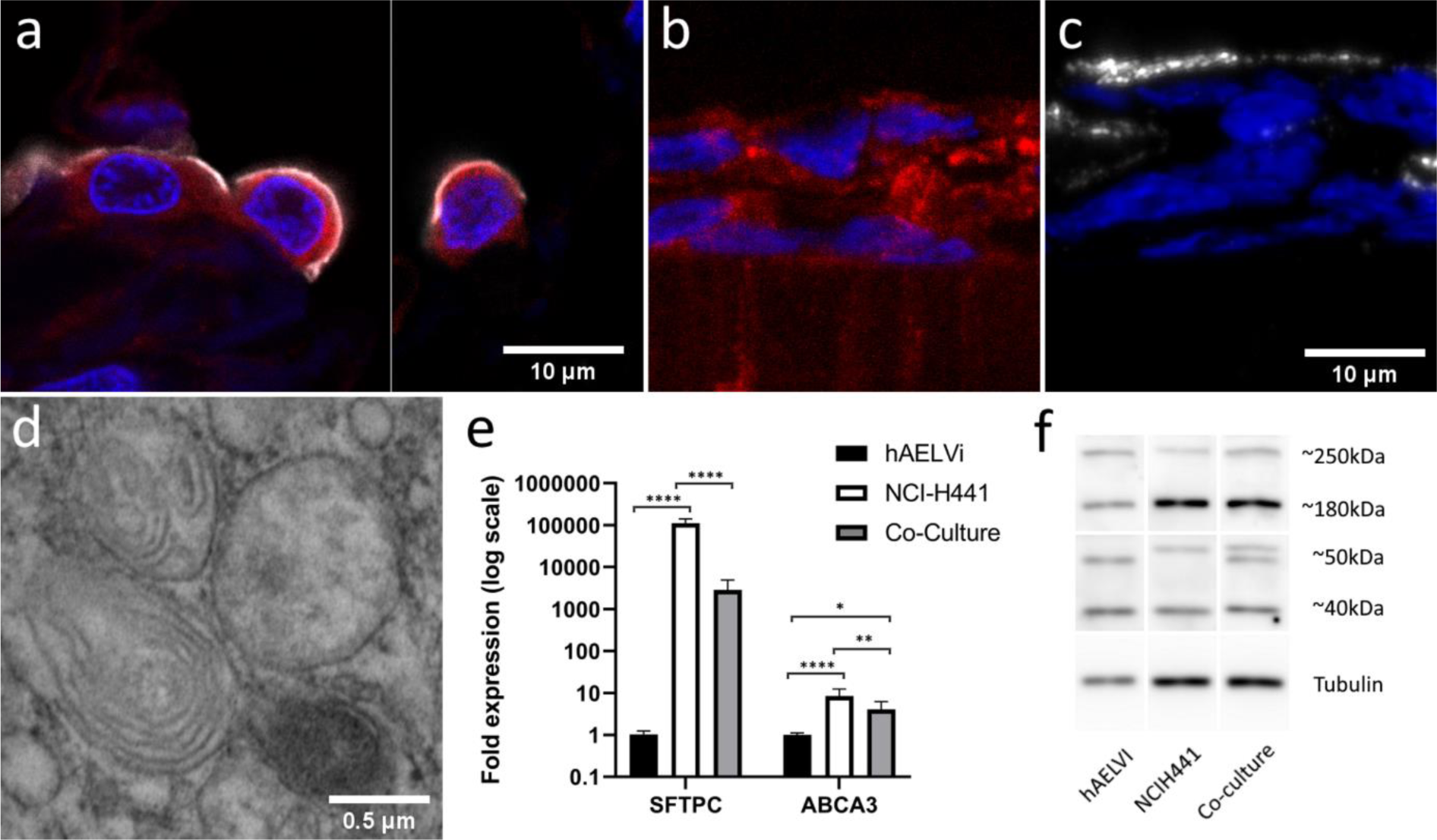
NCI-H441 cells and co-cultures produce surfactant. Immunofluorescent labelling of the lamellar body component ABCA3 (red) in normal human tissue samples (a) and in NCI-H441 ALI cultures (b). HTII-280, another type-II specific antigen (white) is present in the apical surface of type-II cells in normal human tissue (a) and in NCI-H441 (c). Electron micrograph demonstrating the presence of lamellar bodies beneath the surface of NCI-H441 cells grown in ALI conditions (d). RTqPCR reveals expression of both surfactant protein C and ABCA3 (e), and western blotting with an anti-ABCA3 polyclonal antibody reveals post-translational processing of ABCA3 (f).

**Fig 7:**
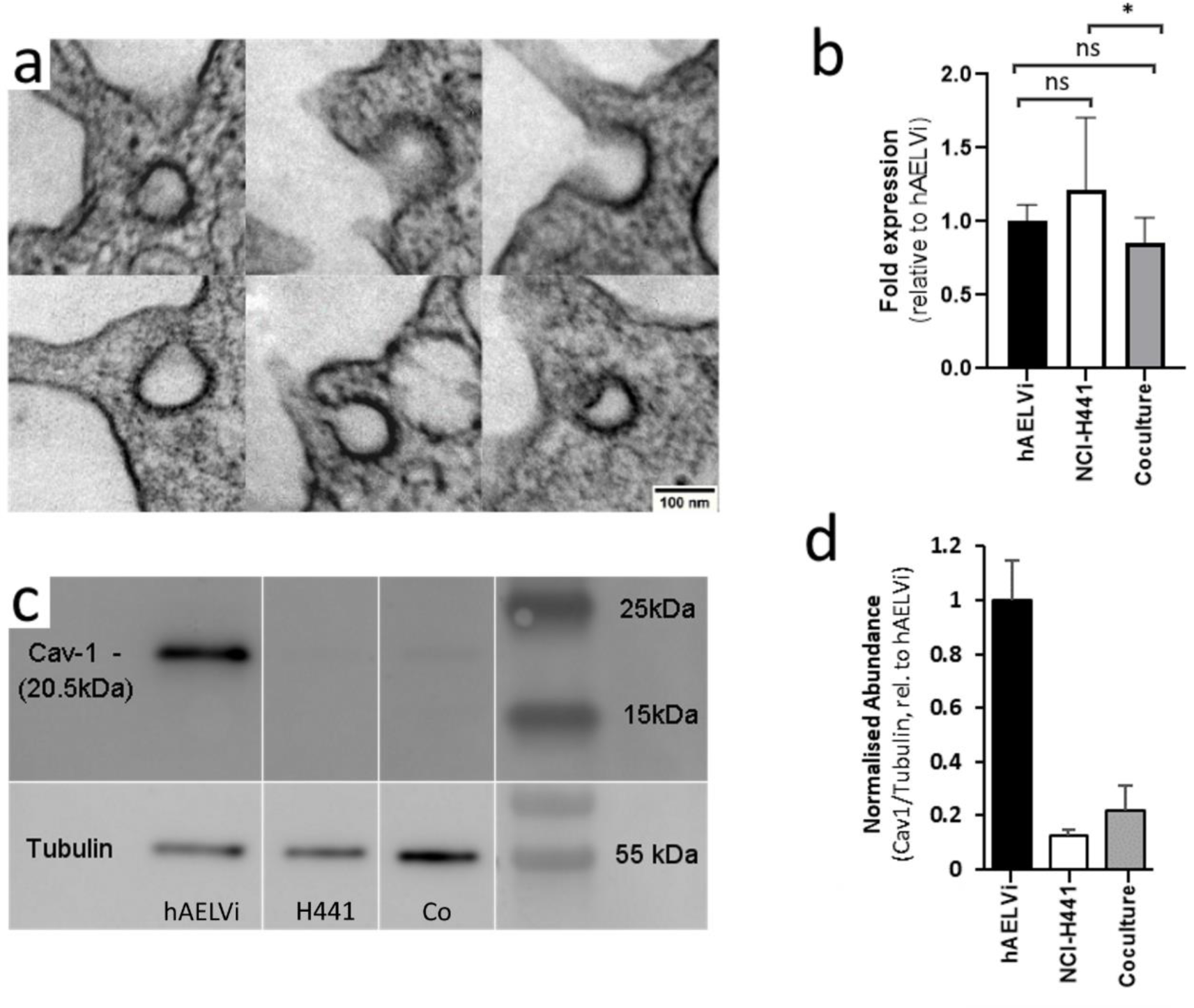
Caveolae are observed in hAELVi, but not in NCI-H441. Single cultures of NCI-H441 and hAELVi, and co-cultures consisting of equal proportions of both were seeded on permeable supports, cultured for two weeks at ALI and harvested. A portion of each membrane was fixed for TEM, and the remaining cells were lysed for extraction of RNA and protein. Structures bearing the ultrastructural features of caveolae were observed in some cells in co-culture (a). No caveoli were observed in NCI-H441 and hAELVi singly. Differences in CAV1 gene expression were not observed (b), but expression of Caveolin-1 protein is substantially higher in hAELVi than in NCIH-441 (c and d). n=3 independent experiments, each containing minimum 3 technical replicates.

**Fig S1. Intracellular void from NCI-H441 culture**

**Fig S2. Modelling the relationship between TEER and Lucifer yellow translocation.** Data were collected from cultures of both hAELVi and NCI-H441 on permeable supports throughout the course of the first two weeks of their development. The graph above shows observed data (red points) of lucifer yellow translocation vs TEER, while the black dotted line shows a function based on the TEER (fTEER) which closely models lucifer yellow clearance. The Solver plugin for excel was used to optimise the offset, scale, and curvature of fTEER to maximise R2 (R2 = 0.98).

**Figure S3. Photograph of original western blot membranes.** Each gel contains three replicates of each condition. Experiments were further repeated three times, consistently reproducing these results. The ladder in the left-hand lane is PageRuler™ (Thermo)

