## Supplementary material for "Co-culture of type I and type II pneumocytes as a model of alveolar epithelium": 20210802_SI-BioRxiv-OB: 20210802_SI_BioRxiv_OB.docx

**Fig S1. Intracellular void from NCI-H441 culture**

**Fig S2. Modelling the relationship between TEER and Lucifer yellow translocation.** Data were collected from cultures of both hAELVi and NCI-H441 on permeable supports throughout the course of the first two weeks of their development. The graph above shows observed data (red points) of lucifer yellow translocation vs TEER, while the black dotted line shows a function based on the TEER (fTEER) which closely models lucifer yellow clearance. The Solver plugin for excel was used to optimise the offset, scale, and curvature of fTEER to maximise R2 (R2 = 0.98).

**Figure S3. Photograph of original western blot membranes.** Each gel contains three replicates of each condition. Experiments were further repeated three times, consistently reproducing these results. The ladder in the left-hand lane is PageRuler™ (Thermo)

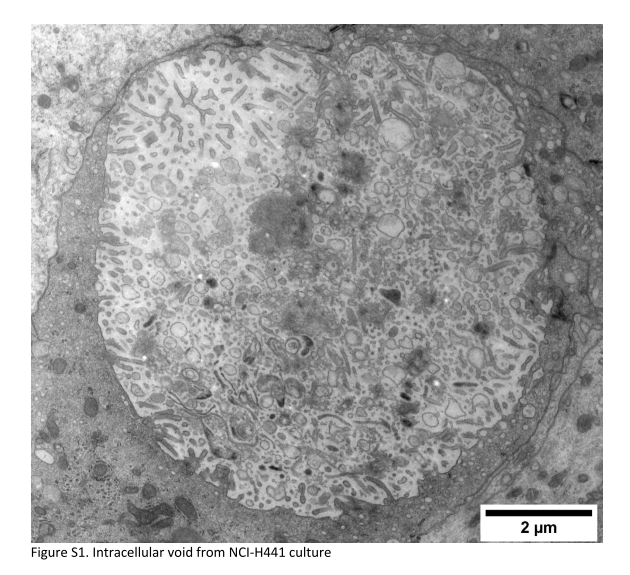

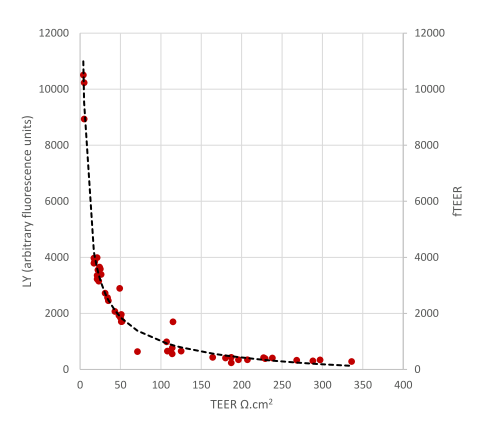

**Fig S2. Modelling the relationship between TEER and Lucifer yellow translocation.**

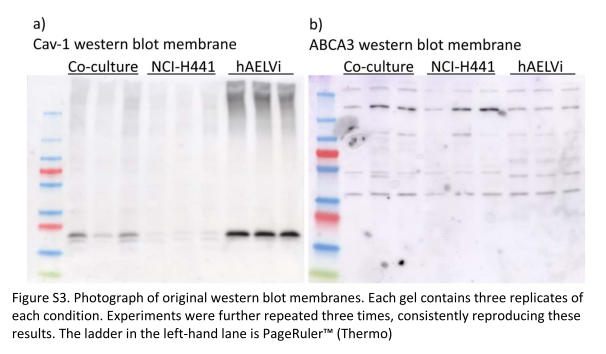
